# *In Silico* Determined Properties of Designed Superoxide Dismutase-1 Mutants Predict ALS-like Phenotypes *In Vitro* and *In Vivo*

**DOI:** 10.1101/474585

**Authors:** Michèle G. DuVal, Luke McAlary, Mona Habibi, Pranav Garg, Mine Sher, Neil R. Cashman, W. Ted Allison, Steven S. Plotkin

## Abstract

The underlying physical causes of SOD1-related ALS are still not well-understood. We address this problem here by computationally designing two de novo mutants, A89R and K128N, which were predicted theoretically to be either significantly destabilizing or stabilizing respectively. We subjected these *in silico* designed mutants to a series of experimental tests, including *in vitro* measures of thermodynamic stability, cell-based aggregation and toxicity assays, and an *in vivo* developmental model of zebrafish motor neuron axonopathy. The experimental tests validated the theoretical predictions: A89R is an unstable, highly-deleterious mutant, and K128N is a stable, non-toxic mutant. Moreover, K128N is predicted computationally to form an unusually stable heterodimer with the familial ALS mutant A4V. Consistent with this prediction, co-injection of K128N and A4V into zebrafish shows profound rescue of motor neuron pathology. The demonstrated success of these first principles calculations to predict the physical properties of SOD1 mutants holds promise for rationally designed therapies to counter the progression of ALS.

**Significance:** Mutations in the protein superoxide dismutase cause ALS, and many of these mutants have decreased folding stability. We sought to pursue this thread using a synthetic biology approach, where we designed two *de novo* mutations, one stabilizing and one destabilizing, as predicted using computational molecular dynamics simulations. We then tested these mutants using *in vitro*, cell-based, and *in vivo* zebrafish models. We found that the unstable mutant was toxic, and induced a severe ALS phenotype in zebrafish; the predicted stable mutant, on the other hand, behaved even better than WT. In fact, it was able to rescue the ALS phenotype caused by mutant SOD1. We propose a mechanism for this rescue, which may provide an avenue for therapeutic intervention.

---

Amyotrophic lateral sclerosis (ALS) is a fatal neurodegenerative disease characterized by progressive loss of motor neurons in the motor cortex, brainstem and spinal cord, with a lifetime risk of about 1:400(1). Most cases of ALS cases are sporadic with no consistently identified genetic mutation, however, a significant fraction of familial ALS (fALS) cases are due mutations in Cu,Zn superoxide dismutase (SOD1)(2-4). The ubiquity of disease-associated mutations (currently over 160(5)) throughout the primary sequence of SOD1 suggests a physico-chemical origin for SOD1-related fALS.

When natively folded, SOD1 is a 32 kDa homodimer that catalyzes the dismutation of oxygen radicals to less harmful molecular oxygen and hydrogen peroxide(4). A properly folded monomer binds one zinc and one copper atom, and contains an intramolecular disulfide bond, all of which are important for the stability of the protein(4). Loss of any of these factors increases the likelihood of SOD1 misfolding into a toxic disease state due to a decrease in stability and invariable formation of toxic aggregates(6). Indeed, SOD1-related fALS mutations are reported to decrease the stability of dimer formation(7, 8), monomer folding(9, 10), and metal binding(11). The variable effects of different mutations on these biophysical components of SOD1 stability and aggregation have been suggested to be responsible for the variable patient survival times in SOD1-fALS(10, 12-14).

In light of this evidence, we were inspired to consider if a *de novo* non-fALS SOD1 mutation could be rationally designed to present phenotypes similar to SOD1-fALS mutants in commonly used experimental SOD1-related fALS models. Conversely, we asked whether a mutant could be designed *de novo* that would further stabilize SOD1 and render it more resistant to misfolding pathology. To this end, we identified two novel mutants, A89R and K128N, which were predicted to be highly destabilizing or stabilizing, respectively. We then assessed the effects of these mutations on protein stability and toxicity using a combination of *in silico* alchemical modelling, *in vitro* protein assays, and mammalian cell cultures, as well as an *in vivo* zebrafish model. We find that the highly destabilizing A89R mutant consistently shows phenotypes similar or worse than the most destabilizing fALS mutations, whereas K128N is consistently wild type-like, as predicted. Furthermore, we predict *in Silico* and show *in Vivo* that K128N can abrogate motor neuron morphology by heterodimer rescue in a zebrafish model when co-expressed with the toxic fALS mutant A4V.

## Results

### Comprehensive residue scans of SOD1 reveal destabilizing and stabilizing mutations

Arginine is overrepresented in observed SOD1-related fALS mutants compared to its expected frequency due to evolutionary mutation (Fig. 1a,b). We thus performed a computational mutation scan of SOD1 by replacing each residue side chain with that of arginine, and obtained estimates of the change in thermodynamic stability for monomers and dimers using the Eris stability prediction software(15). In general, most mutants in this scan were predicted to be destabilizing (Fig. 1c,d), yielding residue A89 as significantly destabilized. A89 is known to have two ALS-associated mutations, A89T and A89V. Other residues were predicted by this method to be even further destabilizing (Fig. 1c,d), however, we focused on A89 because it is not involved in coordinating metals, not in the dimer interface, and is located at the edge rather than center of a β-strand (Fig. 2a), making its expected impact less obvious. Finally, webserver methods are approximate, so we did not rely strictly on the resultant rankings, but rather used these as a starting point for further analysis.

**Fig. 1.**
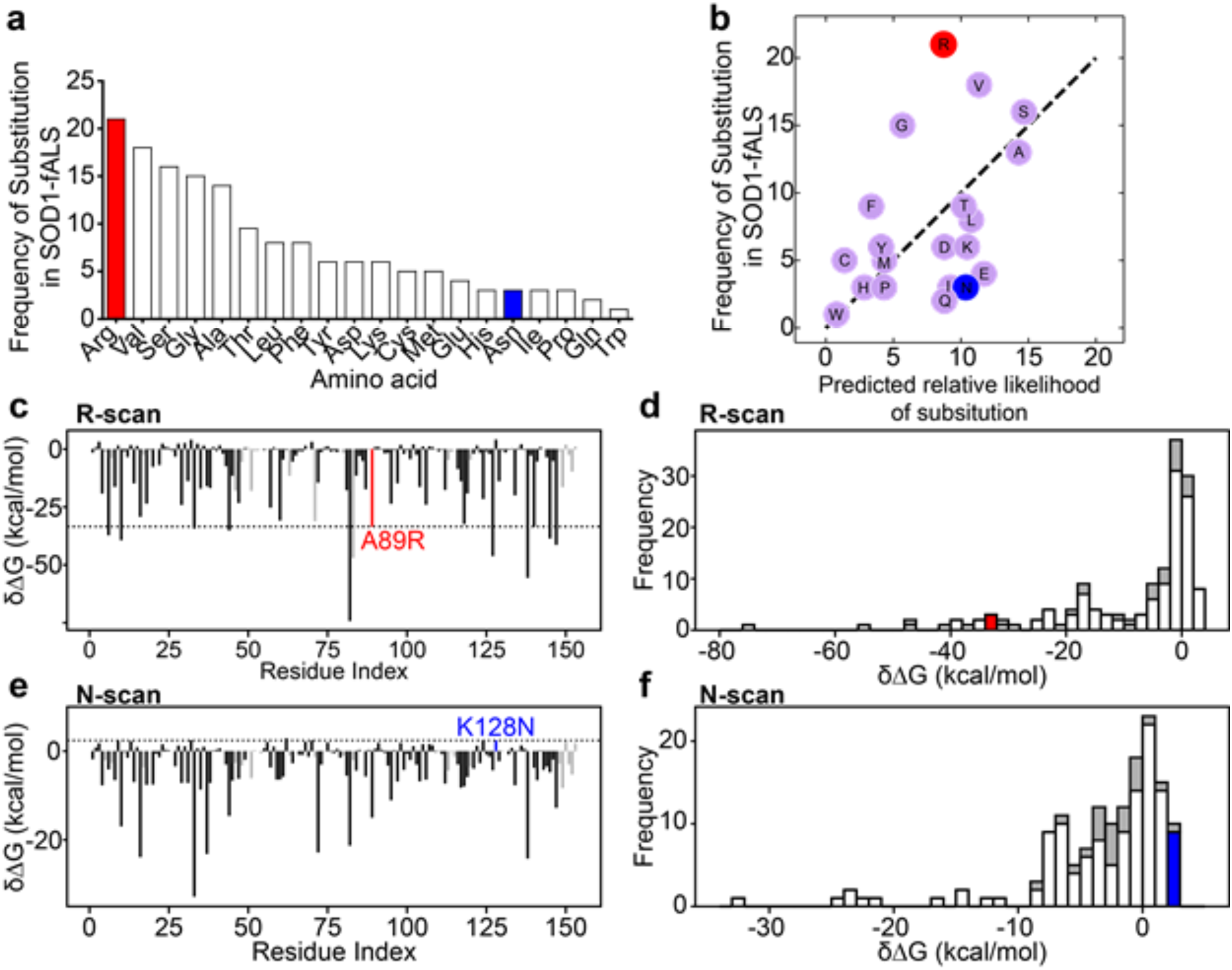
Computational scans support anomalous stability changes for A89R and K128N. **(a)** Amino acids after mutation were gathered from ALSoD showing that Arg substitutions are the most prevalent ALS-causative mutation in SOD1. **(b)** Scatter plot of the frequency of ALS mutations plotted against the predicted likelihood of evolutionary amino acid substitutions from the Kosiol model (see methods). Dashed line has slope equal to unity for comparison. Arg is overrepresented and Asn is underrepresented compared to the evolutionary frequencies of these substitutions. **(c,e)** 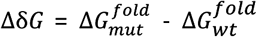 of SOD1 monomers, plotted against residue index for Arg and Asn amino acid scans. Red and blue colored bars indicate the mutants A89R and K128N respectively. Grey bars correspond to residues either in the dimer interface, or those which coordinate metals. **(d,f)** Frequency distribution of δΔG for the **(d)** Arg scan and **(f)** Asn scan with A89R in red and K128N in blue. Grey regions correspond to dimer interface/metal-binding residues.

**Fig. 2.**
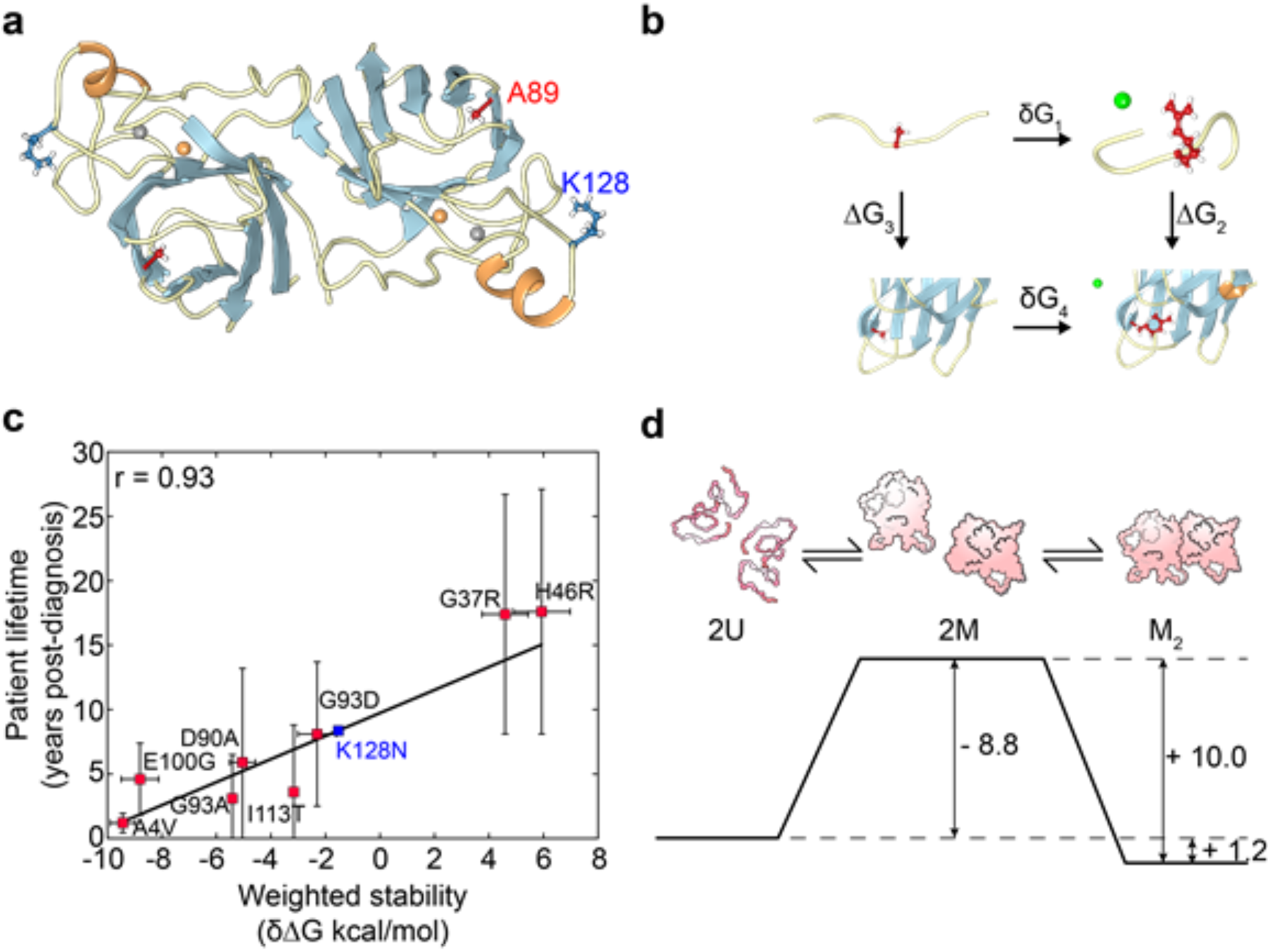
Weighted protein stabilities from alchemical modelling predicts patient survival. **(a)** Native homodimer structure of SOD1 (PDB 1L3N(25)), showing the locations of A89R (red) and K128N (blue). **(b)** Thermodynamic cycle for folding (ΔG_2_, ΔG_3_) and alchemical changes (δG_1_, δG_4_) of a SOD1 monomer alchemical change A89R (red) with chloride counterion (green sphere). **(c)** Patient lifetime correlates with computationally-determined weighted stability of fALS-associated SOD1 mutants (r = 0.93, p = 0.00088). **(d)** Energy landscape of A4V SOD1 to form a homodimer; values are taken from the alchemical free energy calculations along with WT SOD1 numbers from Lindberg *et al* (2005)^10^. Numbers are in kcal/mol.

We also sought to predict beneficial mutants, potentially with greater stability than human WT SOD1. Asparagine substitutions are generally rare among ALS-associated SOD1 mutants compared to their expected evolutionary frequency (Fig. 1a,b)(5), and include only D96N, D101N, and S134N, whereas there are 21 Arg substitutions. We thus implemented an asparagine scan, which yielded K128N as one of the most stabilizing mutants (Fig. 1 e,f).

### Calculating monomer, homodimer, and heterodimer stabilities using alchemical free energy calculations

To more accurately assess the effects of the novel mutants on SOD1 monomer and dimer stability, a systematic approach was taken by using alchemical free energy calculations(16-18). We measured the stability change due to mutation in SOD1 monomers, homodimers, and mutant-WT heterodimers. Apo, disulfide bonded (E,E(SS)) SOD1 monomers or dimers were subjected to an alchemical transformation that introduced a mutant residue in place of a given WT residue, and the change in free energy, mutant to WT, is then calculated.

The effect of a mutation on the folding stability of a monomer was calculated by making alchemical free energy changes on the unfolded and folded states, and using a thermodynamic cycle to find the difference in folding stability, mutant to WT (Fig. 2b). Previous implementations have used a single isolated amino acid to model the unfolded state(17); here, the unfolded monomer was modelled by the equilibrium ensemble of a 7 native amino acid fragment flanked by two glycine residues, with the mutated residue in the middle of the primary sequence. The equilibrium ensemble of the E,E(SS) monomer is used to model the native state. The change in unfolding free energy upon mutation, δ∆*G*_*unf*_ (Fig. 2b), is given by:

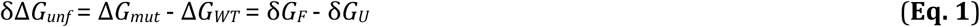

where δ*GX* is the alchemical free energy change of a mutation in either the folded (*F*) or unfolded (*U*) state. Stabilizing (destabilizing) mutations will have a positive (negative) unfolding free energy change. Similarly, the change of dimer stability upon mutation is found alchemically by again using Equation 1, where now δ*G*_*F*_ is the alchemical free energy change in the E,E(SS) native dimer, and δ*G*_*U*_ is 2× the free energy change in a folded monomer. (SI Appendix Fig. S1a).

Based on thermodynamic alchemy calculations, the monomer unfolding free energy for A89R is far less than that of the WT monomer, δ∆*G*_*unf*_ ≈ −31 ± 0.6 kcal/mol. Because ∆*G*_*unf*_ ≈ 4.5 ± 0.1 kcal/mol E,E(SS) for WT SOD1(19), this suggests that with high likelihood, A89R is unfolded under standard conditions. The alchemy number was consistent with the Eris prediction of −33 ± 12 kcal/mol (Fig. 1b). On the other hand, even though the Eris webserver method predicted K128N to be 2.4 ± 0.6 kcal/mol *more* stable than WT SOD1, alchemical calculations predicted K128N as 2.0 ± 0.4 kcal/mol *less* stable than WT (Table 1). Generally the two methods show only modest comparability (SI Appendix Fig. S1c,d).

**Table 1:** Alchemical Free energy changes for novel and ALS-associated mutants.

| SOD1 Mutant | Monomer Unfolding $\delta\Delta G^\S$ | Homodimer Binding $\delta\Delta G$ | Mut-WT Binding $\delta\Delta G$ | Mut-WT Mixing $\delta\Delta G$ |
| --- | --- | --- | --- | --- |
| A4V | $-8.9 \pm 0.2$ | $-0.2 \pm 0.5$ | $-1.4 \pm 0.2$ | $-2.7 \pm 0.7$ |
| G37R | $-0.2 \pm 0.4$ | $8.7 \pm 0.8$ | $-0.70 \pm 0.6$ | $-10.0 \pm 1.4$ |
| H46R | $3.8 \pm 0.4$ | $-3.1 \pm 1.2$ | $12.4 \pm 0.7$ | $28 \pm 2$ |
| A89R | $-31 \pm 0.6$ | $-24 \pm 2.0$ | $1.1 \pm 1.0$ | $26 \pm 3$ |
| D90A | $-1.6 \pm 0.2$ | $-5.1 \pm 0.5$ | $-1.7 \pm 0.3$ | $1.7 \pm 0.7$ |
| G93A | $-3.3 \pm 0.1$ | $-2.9 \pm 0.2$ | $-1.7 \pm 0.1$ | $-0.5 \pm 0.2$ |
| G93D | $-1.3 \pm 0.3$ | $1.7 \pm 0.7$ | $-6.2 \pm 0.5$ | $-14.1 \pm 1.3$ |
| E100G | $-4.9 \pm 0.3$ | $-5.2 \pm 0.7$ | $-3.1 \pm 0.4$ | $-1.0 \pm 1.1$ |
| I113T | $-3.0 \pm 0.1$ | $-0.1 \pm 0.2$ | $-0.4 \pm 0.1$ | $-0.8 \pm 0.2$ |
| K128N | $-2.0 \pm 0.4$ | $-1.8 \pm 0.9$ | $4.7 \pm 0.4$ | $11 \pm 1$ |
| A4V<sup>SH</sup> | $-6.0 \pm 0.1$ | $4.6 \pm 0.5$ | $5.3 \pm 0.3$ | $6.1 \pm 0.8$ |
| A4V<sup>SH</sup>·K128N<sup>SH</sup> | -- | -- | $14.3 \pm 0.3^*$ | $29 \pm 1^\ddagger$ |

<sup>§</sup>All energies are in kcal/mol.
\*Relative to E,E(SH) WT-WT homodimer.
<sup>‡</sup>Corresponding to the process $A4V \cdot A4V + K128N \cdot K128N \rightarrow 2 A4V \cdot K128N$

Similarly, alchemy calculations predict the A89R homodimer is far more unstable than the WT homodimer (δ∆*G*_*dim*_ ≈ −24 ± 2.0 kcal/mol). The K128N homodimer is predicted as slightly less stable (δ∆*G*_*dim*_ ≈ −1.8 ± 0.9 kcal/mol) than the WT (Table 1).

The stability change of the heterodimer upon mutation was also investigated by thermodynamic alchemy. Only one of the two monomers is mutated in this assay. The K128N·WT heterodimer is *more* stable than the WT·WT homodimer, with free energy change (δ∆*G*_*hetdim*_ ≈ 4.7 ± 0.4 kcal/mol).

Interestingly, the A89R·WT heterodimer is also marginally more stable than the WT·WT homodimer, with free energy change (δ∆*G*_*hetdim*_ ≈ 1.1 ± 1.0 kcal/mol, see Table 1).

To determine the *in vitro* properties of these mutants, we first monitored the proportion of SOD1 partitioning to insoluble inclusion bodies when expressed in bacteria in the presence/absence of Cu and/or Zn. We found K128N to be as soluble as WT protein for all metal ion states (SI Appendix Fig. S2a,b). Solubility was generally enhanced by metal cofactors for WT, K128N, and A4V. On the other hand, A89R partitioned almost exclusively into insoluble inclusion bodies, regardless of the presence of Cu and/or Zn (SI Appendix Fig. S2a,b). This extreme insolubility prevented purification of A89R and further *in vitro* characterization.

Expressed SOD1 was purified to obtain thermal stabilities using differential scanning fluorimetry(20). The melting temperatures we obtained for a set of ALS-associated mutants correlated strongly with those previously reported in the literature from differential scanning calorimetry(21) (r=0.95, p=0.0993) (SI Appendix Table2 and SI Appendix Fig. S1h). The melting temperature obtained for K128N (50.4 ± 0.2 °C) was slightly lower than that of WT (51.5 ± 0.3 °C) but greater than all other ALS-associated mutants except for H46R (51.7 ± 0.3 °C).

We further tested the accuracy of the alchemy calculations by correlating with published experimental data(10, 22) for both apo monomer folding stability, and apo dimer binding stability. The calculated monomer unfolding free energies determined here correlated reasonably well with those determined via chemical denaturation (r = 0.73, p = 0.0953) (SI Appendix Fig. S1e), comparable to the correlation between separately-determined experimental values(10, 22) (r = 0.88, p = 0.0519) (SI Appendix Fig. S1f). However, we found no significant correlation between alchemical dimer binding free energies and experimental values (r = 0.60, p = 0.2883) (SI Appendix Fig. S1g). This may be due to the coupled effects of unfolding and unbinding in experiments, or possibly due to long-time relaxations of isolated monomers not captured in simulations; The alchemical method focuses purely on the process of monomerization of the dimer.

As an interesting application of the alchemy method, we asked whether the calculated free energy changes are predictive of patient survival times for known familial ALS-associated mutants(13). Stability changes for monomer unfolding, homodimer binding, and heterodimer binding, are thus calculated for 8 known fALS-associated mutants (A4V, D90A, E100G, G37R, G93A, G93D, H46R,I113T), and the above three thermodynamic quantities are added with two variable relative weights, α and β, to obtain a weighted stability:

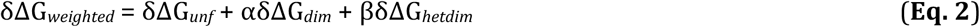

The relative weights α and β are then optimized for best correlation to the corresponding data for the mean patient survival-time from diagnosis for each SOD1 mutant using previously published work(13) (Fig. 2c). The computational data and the patient data show strong correlation (r= 0.93, p= 0.00088). The optimal parameters are (α,β) = (+0.57,+0.32). The fact that α and β are positive implies that all binding free energies are protective against disease. The fact that the magnitude of α and β are both less than the weight of the unfolding free energy implies a ranking of relative importance of each of these processes, with monomer stability being most important.

Strict extrapolation of the best fit between lifetimes and weighted correlation for the A89R novel mutant would register a negative survival time of −30 years; the large change of A89R monomer folding stability indicates that the monomer is likely unfolded, so we might expect a non-linear turn-over in this highly-destabilizing regime, resulting in small positive survival times. In any event, A89R is predicted to be a rapidly progressing mutant. The weighted stability predicts a mean lifetime for K128N of 8.4 years (Fig. 2c). Even more intriguingly, the weighted stability predicts a lifetime after diagnosis for WT SOD1 of 9.7 years, and longer lifetimes than WT for the mutants G37R and H46R. Other factors are certainly at play in governing the initial onset of the disease, which we have not attempted to predict here.

### Heterodimer rescue

Previous in-cell measurements of SOD1 homo-and hetero-dimer concentration using FRET-based constructs indicate appreciable heterodimer concentration across mutants(23). Heterodimerization has been suggested as correlating more strongly with ALS disease progression(12) when some mutants (G37R) were removed from the analysis. On the other hand, several tethered Mutant·WT heterodimer constructs have been shown to be more stable and soluble than Mutant·Mutant homodimers(24).

From the alchemical free energy calculations of mutant·WT heterodimer stability, we can define the mixing free energy (Table 1) as the free energy change corresponding to the process of forming two heterodimers from a mutant·mutant homodimer and WT·WT homodimer: i.e. WT·WT+Mut·Mut⇌2Mut·WT. This gives 2δ∆*G*_*hetdim*_ - δ∆*G*_*dim*_ for the mixing free energy change. For K128N, this process is strongly driven forward by the stability of the heterodimer by about 11 kcal/mol. For A89R, this process is strongly driven forward by the instability of the A89R homodimer. However, this instability of −24 kcal/mol is sufficiently larger in magnitude than the stability of the E,E(SS) WT homodimer (10.2 kcal/mol)(10), so that the homodimer is unlikely to be formed at all. Similarly, as mentioned above, the instability in ∆*G*_*unf*_ for A89R (−31 kcal/mol) is far larger than ∆*G*_*unf*_ for WT E,E(SS) monomer (4.5 kcal/mol) so that the monomer is likely unfolded. One can ask then whether WT·Mut heterodimer formation can “rescue” the folding of A89R. This will happen only if the free energy of the heterodimer is lower than the unfolded free energy. However, from Table 1 and the above WT numbers, ∆*G*_*unf*_ + ∆*G*_*hetdim*_ = −26.5 + 11.3 = −15.2 kcal/mol, so even assuming availability of WT monomer, heterodimer formation is an uphill process, and the thermodynamically-stable state is the unfolded monomer.

Using the alchemical numbers from Table 1 and WT stabilities, along with the uncertainties for both, A4V shows almost zero stability in the homodimer state: ∆*G* for the process *2U→2M→M_2_*, where *M_2_* indicates the homodimer, is 2∆*G*_*unf*_ + ∆*G*_*dim*_ = 1.2 ± 0.9 kcal/mol (Fig. 2d). Heterodimer mixing is unfavorable (Table 1), so the marginally-stable homodimer and the unfolded monomer are the primarily-occupied on-pathway species.

### A89R is prone to form inclusions and is toxic to motor neurons, while K128N behaves similar to WT

A hallmark of SOD1-related fALS is the deposition of mutant SOD1 into intracellular proteinaceous inclusions in patients(2). We assessed the effect of the novel A89R and K128N mutants, as well as other ALS-associated mutants, on inclusion formation in cultured U2OS cells using EGFP-tagged SOD1. We used assisted machine learning with a random forest classifier(26) to identify cells containing inclusions (Fig. 3a,b and SI Appendix Fig. S3,S4).

**Fig. 3.**
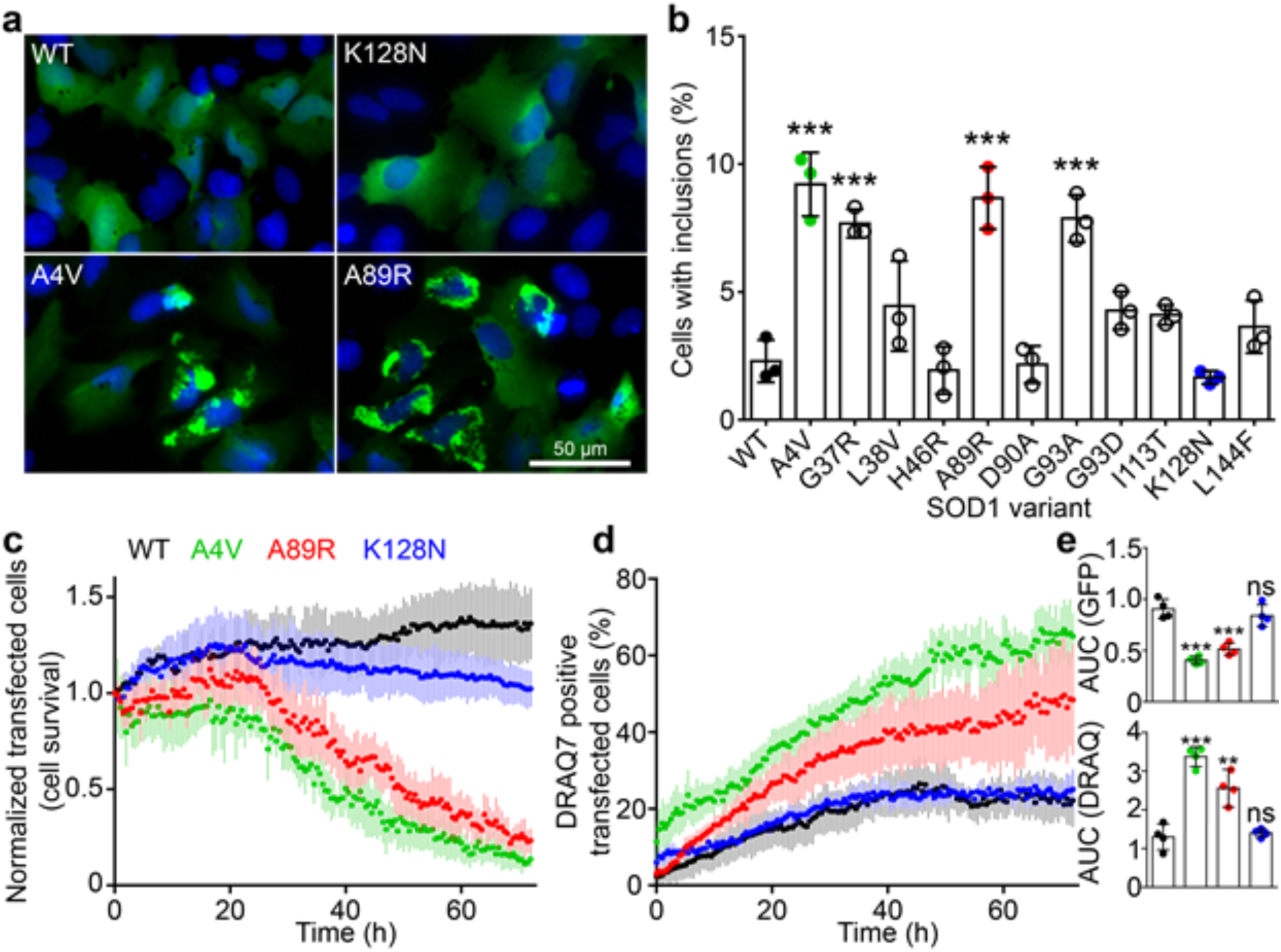
A89R and A4V SOD1 readily form inclusions when expressed in cells, while K128N and WT form few inclusions. **(a)** Representative images of transfected U2OS cells with nuclear Hoescht 33342 (blue) and SOD1-EGFP constructs (green); brighter punctate staining are inclusions. Scale bar is 50 µm. **(b)** Quantification of the number of U2OS cells with inclusions by machine learning showed significant increases (p < 0.01) in the percentage of inclusion-containing cells for A4V, G37R, A89R, and G93A compared to WT. **(c)** Quantification of the number of surviving NSC-34 cells *vs.* time relative to the starting number, during a live cell assay. **(d)** Quantification of the percentage of DRAQ7 positive transfected cells *vs*. time. **(e)** Area under the curve for GFP counts and for DRAQ. Data for inclusion formation is n = 3 biological replicates where error bars represent SD of the mean. Live-cell toxicity assays were 4 biological replicates where error bars represent SEM and statistical significance of the AUC was determined using one-way ANOVA with Dunnet’s post-test with WT as control (*** *p* < 0.01, ** *p* < 0.05, *ns = not significant*).

We found that cells expressing A4V, G37R, or G93A were significantly more likely to contain inclusions, while WT, H46R, and D90A formed very few inclusions, in agreement with previously published work (Fig. 3 a,b)(27). Mutants L38V, G93D, I113T, and L144F all trended towards a greater number of inclusion-containing cells, however, the observed difference from WT was not found to be statistically significant (Fig. 3 b). A89R showed inclusion formation on-par with the highly aggregation-prone and toxic A4V-fALS mutant, whereas K128N showed the lowest mean fraction of inclusions, comparable to WT (Fig. 3 a,b).

We further assessed the cellular toxicity of our SOD1-EGFP mutants by transfecting them into the mouse motor neuron-like cell line NSC-34, and examined both the GFP positive cell population and plasma membrane integrity(28) *vs*. time. The mutants A4V and A89R showed significantly attenuated levels of GFP positive cells *vs*. time (Fig. 3c,e), with A4V in agreement with previous work(27). On the other hand, K128N showed continued proliferation and no significant difference from WT in this assay (Fig. 3c,e). Cells expressing A4V or A89R were significantly more likely to uptake DRAQ7 dye (~3× and 2.5× respectively) (Fig. 3c,e). In contrast, WT and K128N showed comparable levels of DRAQ7 uptake (Fig. 3d,e). This indicates that expression of either A4V or A89R was far more toxic than the expression of either WT or K128N in this model, further supporting the computational predictions and folding stability measurements as predictive of phenotypic outcomes.

### In vivo motor neuron phenotypes by novel SOD1 mutants confirm predicted severities

Having established the effects of the A89R and K128N mutations in cell lines, we next sought to examine the consequences of these mutants in a whole organism system containing motor neurons, by comparing phenotype severity due to mutant SOD1 expression *vs*. WT. We thus expressed SOD1 mutants in a transgenic zebrafish (*D. rerio*), expressing GFP (*Tg[mnx1:GFP]*) in trunk motor neurons and axons (Fig. 4 a,b), and then imaged the axon morphology. Atypical branching patterns in motor neurons as a result of mutant SOD1 expression is an effective *in vivo* system to determine SOD1 pathology(29).

**Fig. 4.**
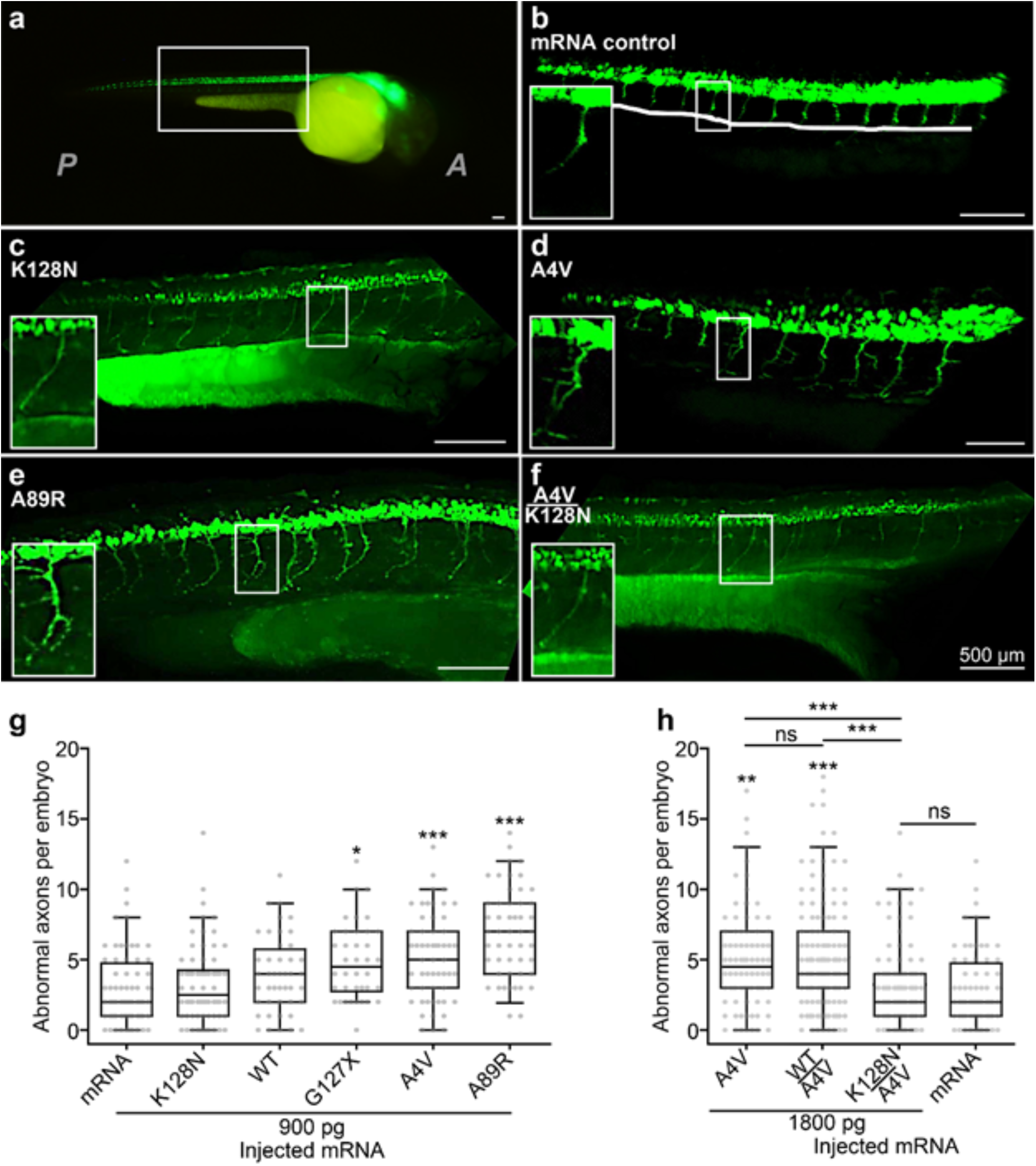
Axonopathy due to mutant SOD1 RNA in zebrafish. **(A-F)** Tg[*mnx1:GFP*] zebrafish embryos were injected with either control mRNA or mRNA encoding SOD1. Scale bars = 100 µm. **(a)** Image of whole embryo illustrating GFP expression in motor neurons (green) in the spinal cord. *A*= Anterior; *P*= Posterior. **(b)** Injection with control mRNA causes little axonopathy or branching above the ventral notochord boundary. Representative image depicting the notochord (white line) with healthy unbranched motor axons. Insets in all images show magnification of a motor axon. Representative images of motor axons of embryos injected with mRNA encoding **(c)** K128N SOD1, **(d)** A4V SOD1, **(e)** A89R SOD1, **(f)** co-injection of A4V and K128N. **(g)** Quantification of the number of abnormal axons per embryo; control mRNA or SOD1 mutant mRNA and amount injected are labelled underneath the x-axis. **(h)** Quantification of axonopathy from fish co-injections at a 1:1 ratio. mRNA control in (g) is included for comparison. For both **g** and **h**, centre line=mean; box limits=upper and lower quartile; whiskers=5% and 95% range. Each treatment group represents at least 39 injected embryos (see Methods for group sample sizes). Statistical significance compared to the control mRNA injected group is represented by asterisks, except asterisks with underlines, which indicate statistical significance of pairwise comparisons. Tests performed were Kruskall-Wallis test with post-hoc Mann-Whitney pairwise comparisons. (ns = not significant, * *p*< 0.05, ** *p*< 0.01, *** *p*< 0.005).

We compared mRNA control injections and five SOD1 mutants (Fig. 4 b-g): WT SOD1, A4V, G127X, and the novel mutants A89R and K128N. A4V and G127X were included in order to compare two known fALS-associated mutants. Other ALS-associated mutants, including G93A and G37R, have been previously assayed(29). Because the above computational and *in vitro* analyses compare the stability and aggregation of SOD1 mutants relative to WT, we evaluated axonopathy of SOD1 mutants relative to axonopathy induced by WT SOD1, in addition to control mRNA injection.

We found that both A4V and the novel mutant A89R caused significantly more axonopathy than mRNA control injection, K128N, or WT SOD1. In fact, the A89R mutant increased axonopathy 30% more on average than the next most toxic SOD1-fALS mutant tested (A4V). The mutant G127X had an intermediate effect less severe than either A4V or A89R (Fig. 4g). Meanwhile, K128N showed axonopathy levels similar to mRNA control and lower than WT (Fig. 4c,g).

The healthy phenotype of K128N compared to WT prompted us to speculate whether K128N could rescue the toxicity of A4V *in vivo*, and if so, whether this might be explained by a direct interaction such as A4V·K128N heterodimer rescue. To assess this computationally, we performed thermodynamic alchemy for two species: disulfide oxidized E,E(SS) SOD1, and disulfide reduced E,E(SH) SOD1. Disulfide reduction has been shown to destabilize the dimer for WT protein(19), however, recent NMR evidence indicates that a native-like homodimer structure, as well as other, non-native dimer structures, are transiently occupied for E,E(SH) SOD1(30, 31). Furthermore, mutant SOD1 can access additional conformations that can lead to possible pathogenic intra- and intermolecular contacts(31). In the absence of further quantification, we assume the E,E(SH) WT homodimer has zero stability relative to two monomers, but the observations from NMR studies(30) suggest the possibility that relatively stable dimers may exist for some mutants in the E,E(SH) state.

The strongest computational evidence that might argue for A4V·K128N phenotypic rescue involves an unusually strong heterodimer between A4V and K128N in the E,E(SH) form: The free energy change, from a K128N homodimer and an A4V homodimer, to two A4V·K128N heterodimers, is: 2δ∆*G(A4V·K128N) - δ∆G(A4V·A4V) - δ∆G(K128N·K128N)* = 29 kcal/mol.

This results in a deep, protective region of the folding and maturation landscape (Fig 5), for which entrenchment would incur a long uphill climb thermodynamically to access those regions where thermostable misfolding and aggregation(32) and/or cellular pathology would take place(33). This large degree of protection appears computationally to be specific to premature A4V, and is not conferred to more mature forms of A4V: The free energy change to form two E,E(SS) A4V·K128N heterodimers from their corresponding homodimers is −5.5 ± 1.3 kcal/mol. The landscape in Fig. 5 illustrates the rescue scenarios for A4V·K128N, along with the landscapes for E,E(SS) WT and A4V·WT heterodimers.

**Fig. 5.**
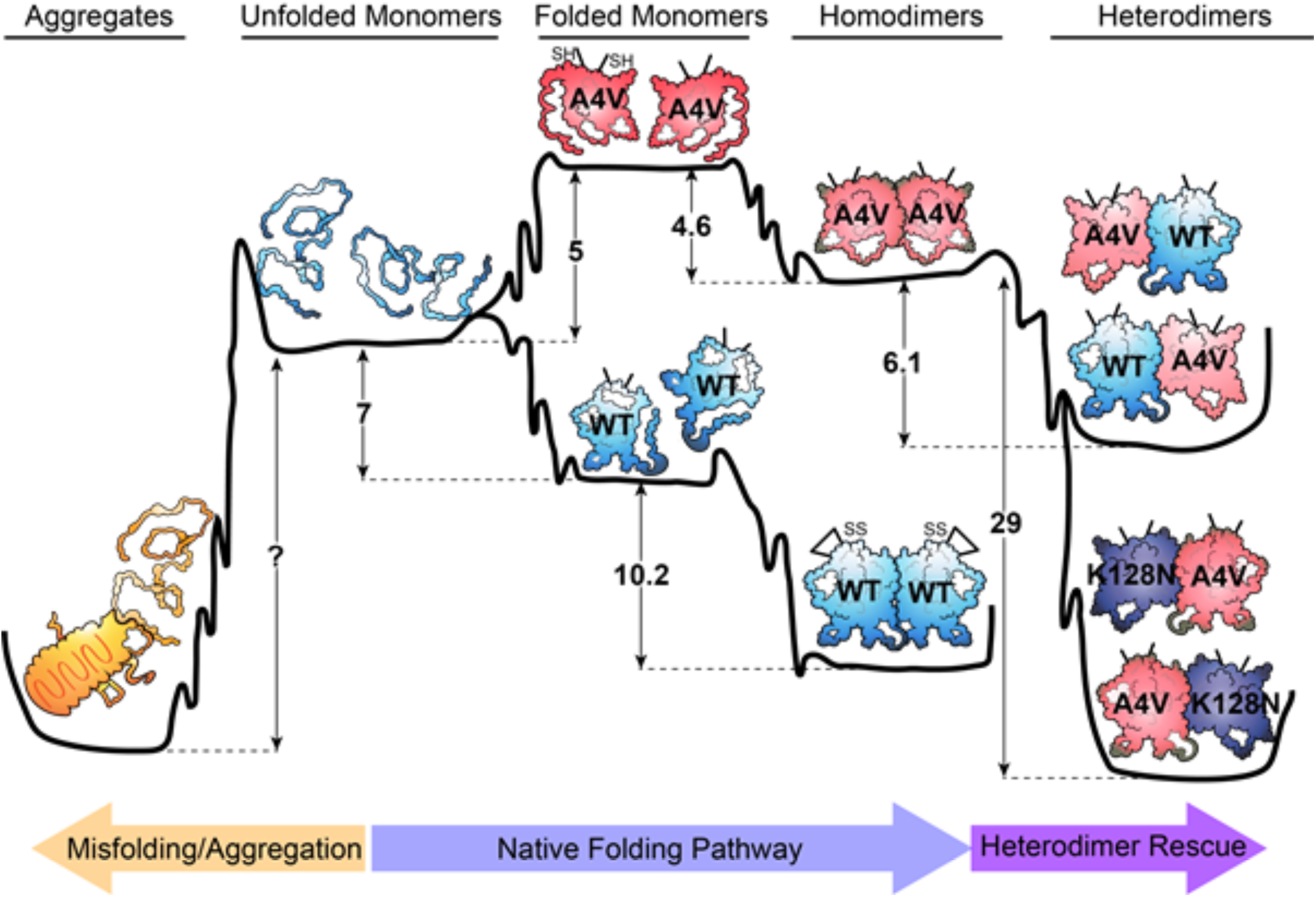
Free energy landscape of K128N-mediated heterodimer rescue of A4V SOD1. Schematic landscape showing the folding and maturation to biologically functional states from unfolded monomers, as well as misfolding to cellularly toxic species (top labels). WT SOD1 (blue), A4V (red), K128N (purple), and aggregates (orange) are all shown, along with corresponding free energy changes (in kcal/mol) between various maturation states. Deeper minima on the right hand side provide more protection from transitions to the deep minimum on the left hand side.

Consistent with these theoretical calculations, dosage-controlled co-injections of K128N/A4V showed significantly less axonopathy than either co-injections of WT/A4V, injections of A4V alone at the total dosage, or injections of A4V at half the dosage (p < 0.005, Fig. 4d,f,h). In fact, co-injections of K128N/A4V showed no significant difference in axonopathy from either the WT (p= 0.06158) or control mRNA (p= 0.6395) half-dosage (900pg) controls (Fig. 4h, Mann-Whitney pairwise comparisons). This suggests a protective interaction imparted by K128N upon A4V, wherein K128N ameliorates the neuronal toxicity caused by A4V. As mentioned above, such a protective niche is not predicted computationally for A4V·WT heterodimers; similarly, WT·A4V co-injections showed significant pathology over controls, and no significant difference from A4V injections alone (Fig. 4h).

## Discussion

In this work we pursued the hypothesis that the thermodynamic properties of SOD1-fALS mutants may be calculated from first principles using computational methods, and that such measurements could be used to predict phenotypic outcomes for *in vitro* and *in vivo* systems. We were thus able to design two *de novo* SOD1 mutants not previously studied, A89R and K128N, and validate their predicted effects in *in vitro* and particularly in an *in vivo* model of SOD1-fALS.

Previous novel SOD1 mutants have been constructed by point mutation, including W32S/W32F(34, 35), C111S(36), T2D(37), and E133Q(38), however, these studies did not use predictions of thermodynamic stability as a condition for their selection. Other studies have analyzed the force-induced mechanical unfolding of a *de novo* designed SOD1 construct^.(39)^, obtained by truncating the metal-binding loops(40). *In silico* methods have also been successfully used to design mutants of β2-microglobulin by changing surface residues to modulate its intrinsic solubility(41).

Here, we found that for the *de novo* mutants A89R and K128N, thermodynamic stability, aggregation propensity, cellular toxicity, and even *in vivo* axonopathy were consistent with the *in silico* predicted effects for each mutant. We also considered several fALS mutants for comparative analysis. To our knowledge, there is no published work which has attempted to computationally predict SOD1 mutants that would be highly pathogenic or potentially protective, and then subsequently validated these predictions experimentally. The prediction of highly pathogenic protein mutants is a powerful tool for delineating mechanisms by which proteins may cause toxicity in neurodegeneration. Our examination of A89R from *in silico* predictions to *in vivo* experiments supports the notion that the folding stability of SOD1 is a significant factor in the ability of a mutant to elicit toxicity, providing a test of the thermodynamic hypothesis using novel SOD1 synthetic biology.

Our initial screens predicted several mutants that had more extreme effects on stability than the mutants we explored here. These mutants include G82R with a predicted δ∆*G* = −74 kcal/mol and P62N with a predicted δ∆*G* = +2.9 kcal/mol. These mutants are in metal binding regions however; we discarded these to avoid conflating the problems of stability and metal binding. A89 and K128 are remote from the metal binding regions in SOD1, but may still affect metal binding allosterically.

It is clear that metals are present in the cell-based and *in vivo* systems, and therefore may be playing a significant role in native protection from pathology(42). Considering this, an interesting prospect would be the *in silico* examination of the metal binding properties of *de novo* SOD1 mutants in order to elucidate the contribution of metal binding to folding stabilities. This non-trivial study would demand force field reparameterization of some key amino acids (e.g. the doubly-deprotonated His63) by quantum chemical methods to properly model copper and zinc coordination(43).

Cell-based and *in vivo* systems also contain endogenous SOD1 which may interact with exogenously expressed SOD1. Here, we have not explored the energetics of heterodimer binding or aggregation-related interactions between mutant human SOD1 and murine or *D. rerio* SOD1 (which have 81% and 76% sequence similarity with human WT respectively). Future studies using suitably tagged SOD1 protein could potentially examine direct interactions *in vivo(23)*.

Our *in silico* and *in vivo* studies suggest that heterodimer rescue provides an additional protective mechanism against pathological misfolding: The K128N·A4V heterodimer appears computationally to be strongly stabilized in the disulfide-reduced state, apparently overcoming any extra loop entropy penalty(19), and correspondingly, K128N co-expression was able to rescue A4V axonopathy. A4V has been observed to be unfolded and disulfide reduced *in vivo*(44). The extra stability imparted by heterodimerization may prevent further loss of native structure in the absence of disulfide bond formation, and may prevent misfolding-associated pathology by sequestration of the precursors to cellularly-toxic species. *In vivo*, Zn ions may stabilize the nascent protein and allow for a stable heterodimer; dimer rescue of SH species by metalation has been observed *in vitro*(45).

A first-principles understanding of the allosteric network of interactions resulting from the *de novo* mutants that we predicted here has the potential to guide basic scientific research as well as rational therapeutic strategies. The exploration of mutants by rationally-guided design can aid the development of “super-proteins” having reduced propensity for misfolding and aggregation as well as enhanced functional and thermodynamic properties, and is an interesting topic of future research.

## Materials and Methods

All simulations were performed with GROMACS-4.6(46), the CHARMM22*(47) force field, with TIP3P water and 150 mM NaCl, solvated in a dodecahedron box. All protein was purified from *E. coli* as described previously(8). Cells were cultured and transfected as previously reported(27). Detailed methods are provided in SI Appendix. Fish care and experimental protocols were approved under the protocol AUP00000077 by the Animal Care and Use Committee: Biosciences at the University of Alberta under the auspices of the Canadian Council on Animal Care.

## Acknowledgements

The authors would like to thank Associate Prof. Justin Yerbury (University of Wollongong, Australia) for access to SOD1 bacterial and mammalian expression plasmids. We also thank Prof. Nikolay Dokholyan (Penn State College of Medicine, USA) for access to the ERIS software executables. S.S.P., W.T.A., and N.R.C. are supported by the Canadian Institutes of Health Research. S.S.P. is also supported by the Alberta Prion Research Institute (APRI) and Compute Canada/Calcul Canada.

## Supplementary Information

**SI Methods and Materials**

### Arginine and alanine scans to predict deleterious and protective mutants

Eris protein stability prediction software (v1.0) was used in fixed-backbone mode with two chains each from Protein Data Bank (PDB) structures 2V0A(5), 2C9V(6), 1HL5(7) (O,Q) and 3ECU(8) (A,B) to estimate the change in free energy of folding upon mutation. The mutants studied included all possible single point mutations to arginine (Arg), asparagine (Asn) and alanine (Ala). In the Ala scan, any alanine residues were mutated to glycine. In the Asn and Arg scans, any Asn or Arg residues were left unchanged. We also ran the same calculation for all possible point mutations at residue 128 (K128*). For any particular mutation, the mean and sample standard deviation of eight values (two chains each from four PDB structures) was used as final output. Approximately 1% of the values were nonsensical (31 out of 3548 calculations failed), and were removed from the analysis; this occurs possibly because the Eris software fails to run correctly with certain combinations of mutation and input structure, perhaps due to unresolvable constraints.

The most destabilizing mutants from the arginine scan, and the most stabilizing mutants from the other three scans were collected as potential candidates for further analysis. We eliminated any mutants involving residues that bind metals, and those in the dimer interface. The set of residues in the dimer interface (5, 7, 17, 50-54, 113-115, 148-153) was obtained by calculating the change in solvent accessible surface area upon dimerization, using the DSSP software(9). Out of the remaining mutants, we chose to use A89R as a promising destabilizing mutant and K128N as a potentially stabilizing one.

### Comparison of SOD1-fALS mutation frequencies against an evolutionary model

The frequencies of the mutant amino acid resulting from the 159 unique exonic substitution mutants listed in ALSoD(10) (as of June 2018) were compared against an empirical codon substitution model, fitted to extant vertebrate sequence data(11). The codon substitution rates were multiplied by their frequencies in the SOD1 exons to find the rate of appearance of each codon. Using the standard codon table, this was used to calculate the rate of appearance of each mutant amino acid from all possible non-synonymous mutations. These rates were rescaled by a multiplicative normalization constant so that when summed over the mutant amino acid identities, they would add to 159. This was done in order to compare against known mutant frequencies (Fig. 1b). The straightforward analysis here does not account for the more subtle effects of variation in mutability across the SOD1 gene, nor the penetrance of the mutations (which affects diagnosis rates).

### SOD1 mutant modeling and free energy change calculations using the alchemy method

The difference in unfolding free energy of apo(SS) SOD1 monomer, mutant to WT, was calculated by thermodynamic alchemy (12-14). Similarly, thermodynamic alchemy was used to calculate the difference in the free energy cost to monomerize the homodimer, mutant to WT, and to monomerize the WT-mutant heterodimer compared to the WT homodimer. PDB ID 2V0A (5) was used as a starting structural model for these calculations.

All simulations were performed with GROMACS-4.6 (15), the CHARMM22* force field(16), with TIP3P water and 150 mM NaCl, solvated in a dodecahedron box. NaCl and KCl have similar effects on protein stability in molecular dynamics simulations, with Na^+^ binding more strongly to proteins than K^+(17)^. The nonequilibrium fast-growth thermodynamic integration method was used to calculate free energy changes due to mutation(13). Hybrid topology files and mutations were generated by the pmx software as described in references (13, 18). Each 7 amino acid peptide system included about 6000 water molecules in box size of 6.4 × 6.4 × 4.5 nm^3^. The systems with single monomer were solvated in a box with size of 7.9 × 7.9 × 5.6 nm^3^ with about 11000 water molecules. The dimer systems were in a box of size 10.0 × 10.0 × 7.1 nm^3^ with about 22100 water molecules.

Each system was equilibrated for 10 ns using a stochastic dynamics integrator at NPT ensemble at T=298 K and constant pressure of 1 atmosphere. The pressure was kept constant by Parrinello-Rahman barostat (19). A cutoff of distance of 1.1 nm and a switching function between 1.0 and 1.1 nm were used for Lennard Jones interactions. Particle-mesh Ewald method (20) was used to calculate electrostatic interactions.

We compared alchemically derived energies of homodimerization to values reported in Lindberg *et al* (2005)(1). In expressions below, U = unfolded monomer, N = native monomer, N_2_ = dimer. Experimental mutational changes in dimer binding free energies δ∆*G_N2-2N_* were obtained by using the formula:

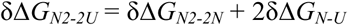

which corrects the formula in ref.(1) by including twice the monomer unfolding free energy in the total free energy change. Our alchemical numbers obtained directly for the monomerization process do not show good agreement with experimental numbers obtained using the above formula by subtracting unfolding free energies from total free energy changes. This may be due to the coupled effects of unfolding and unbinding in experiments, or possibly due to long-time relaxations of isolated monomers not captured in simulations. To obtain unfolding free energy changes for mutant SOD1, the WT monomer unfolding free energy ∆*G*_N-U_ in Lindberg et al (2005)(1) was used as a reference.

### Mutations of amino acids that involve a change in side-chain charge

Several mutations involved a change in charge between the initial and final states, for example mutating Alanine (*q*=0) to Arginine (*q*=+1). These mutations included A89R, D90A, E100G, G37R, G93D, H46R, and K128N. In order to maintain charge neutrality at all the times during fast growth alchemical simulations, counterions need to appear or disappear dynamically and continuously during the alchemical process, with the number and sign of the counterion charges depending on the change in system charge as follows. If after any mutation, the system charge changed by +1 (e.g. for A89R, G37R, H46R, E100G, D90A), then a Cl− ion was grown. If after any mutation, the system charge changed by −1 (e.g. K128N, G93D), we made a Cl− ion disappear. To implement this process, we continuously grow or diminish counterions as follows. In the first step, we gradually turn off all of the partial charges on the residue to be mutated, and calculate the corresponding free energy change ∆*G_1_* as that corresponding to turning off all electrostatic interactions while retaining van der Waals interactions. Next, we mutate the now neutral amino acid to the new topology of the final amino acid, while growing (or diminishing) a dummy particle into a neutral “counterion”; the corresponding free energy change for this process is ∆*G_2_*. Finally, we gradually turn on the charges on all atoms of the mutated residue and counterion and calculate the corresponding free energy change as ∆*G_3_*. The total free energy change for mutations involving a change of charge is thus ∆*G_tot_* = ∆*G_1_* + ∆*G_2_* + ∆*G_3_*. For all mutants where the total charge does not change, the mutation is done all in one step.

### Linear model for patient lifetime prediction

Data for patient lifetime post diagnosis from several studies is collected together in Table 6 of (21). From this table, lifetime data corresponding to deceased patients was used to calculate a mean and sample standard deviation for the mutations of interest (i.e, values such as “>17.2” were excluded). The following expressions were used for this calculation:

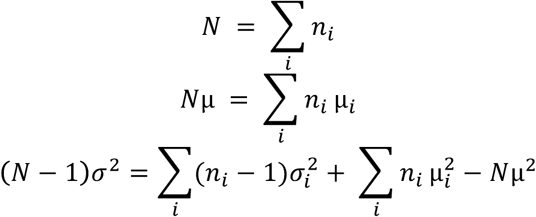

where (*N*, μ, σ^2^) are the sample size, mean and variance of the aggregated data from the various studies for that mutant, and 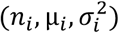 are the sample size, mean and variance of data from the *i*^th^ study of that mutant.

### Plasmids used for Recombinant Protein Purification

Expression vectors encoding SOD1-WT and SOD1-G93A for bacterial expression were a kind gift from Professor Mikael Oliveberg (Stockholm University, Sweden). Expression vectors encoding A4V, G37R, H46R, D90A, G93D, and V148G were kind gifts from A/Prof Justin Yerbury (University of Wollongong, Australia). Expression vectors encoding L38V, A89R, I113T, K128N, and L144F were designed in-house and generated by Genscript (New Jersey, USA).

### Plasmids for Mammalian Transfection

Plasmids for the expression of SOD1 in mammalian cells included pEGFP-N1 encoding C-terminally GFP-tagged SOD1 variants WT, A4V, G37R, H46R, D90A, G93A, G93D, and V148G were kind gifts from Professor Justin Yerbury (University of Wollongong, Australia). Vectors for the expression of pEGFP-SOD1 encoding L38V, A89R, I113T, K128N, and L144F were designed in-house and generated by Genscript (USA).

### Expression and Purification of Recombinant SOD1

Protein expression and purification were performed according to previous methods(22) with modification as follows. SOD1 expression vectors including the gene for yeast copper chaperone were transformed into competent BL21(DE3) *E. coli* via heat-shock. Expression cultures in 1× TB were grown until the OD_600_ was 0.6, at which point ZnSO_4_ and CuSO_4_ were added to 200 µM and 3 mM respectively, and expression was induced by IPTG (0.5 mM) followed by overnight incubation at 20 °C with orbital shaking (300 rpm). Following overnight incubation, cells were collected via centrifugation (6000*g* for 10 min) and resuspended in 50 mM Tris-base (pH 7.4). Cells were then sonicated (25 sec duty cycle) and forced through a 23 gauge needle, with this process repeated five times. Cellular debris was removed via centrifugation (30,000*g* for 20 min) and the supernatant was incubated at 65 °C for 30 min followed by centrifugation to clear any aggregated protein (30,000*g* for 20 min). Ammonium sulphate cuts of 60% (w/v) and 90% (w/v) were performed at 4 °C with moderate stirring, where precipitated material was cleared via centrifugation (30,000*g* for 20 min); the 90% cut was incubated overnight. Following overnight incubation, the 90% cut was pelleted by centrifugation (30,000*g* for 20 min) and the pellet was resuspended in 50 mM Tris, 150 mM NaCl (pH 7.4). Samples were loaded onto a gel filtration column (Hiload 16/60 Superdex 100 PG, GE, USA) equilibrated in 50 mM Tris, 500 mM NaCl (pH 7.4) and eluted at a flow rate of 0.8 mL/min. Fractions containing SOD1 (determined by SDS-PAGE) were pooled and dialysed into 20 mM Tris (pH 8) using SnakeSkin™ dialysis tubing (Thermofisher, USA) with three buffer changes. Dialysed sample was then loaded onto an anion exchange column (HiScreen CaptoQ, GE USA) equilibrated in 20 mM Tris (pH 8) buffer and was eluted across a 0 – 250 mM NaCl gradient. Fractions containing pure SOD1 (determined by SDS-PAGE) were pooled and concentrated using centrifugal concentrators (Millipore, 10 000 MWCO, GE USA). Concentrated samples were flash frozen in liquid nitrogen and stored at −20 °C until use.

### Differential Scanning Fluorimetry

Purified SOD1 was demetallated(23) to generate E,E(SS) SOD1. E,E(SS) SOD1 was dialysed into 10 mM HEPES, 150 mM NaCl (pH 7.4) buffer overnight at 4 °C, flash frozen in liquid nitrogen and stored at-20 °C until use. E,E(SS) SOD1 had its concentration determined via BCA assay (Pierce BCA protein assay kit, Thermofisher, USA) prior to analysis. Protein was plated into a 96-well fast-PCR plate (Thermofisher, USA) at a concentration of 20 µM monomer with 20× SYPRO™ Orange (ThermoFisher, USA), and a 20 µM chicken egg-white lysozyme (Biobasic, Canada) control. A QuantStudio-6 Real-Time PCR machine (ThermoFisher, USA) was used to measure dye fluorescence with a FAM filter-set, controlled by QuantStudio Realtime PCR software (version 1.3 - ThermoFisher, USA). Plates were heated at a rate of 1 °C/min until a final temperature of 95 °C was reached. An in-house python script was written to obtain the melting temperature (T_m_) from the maximum absolute magnitude of the first derivative of the raw data.

### Bacterial Protein Expression Solubility Assay

BL21(DE3) *E. coli* housing expression vectors for SOD1 variants and a yeast copper chaperone were grown in 1× LB supplemented with carbenicillin (100 µg/mL). Starter cultures were used to inoculate main cultures of 1× LB with 1 % glucose (w/v) and carbenicillin (100 µg/mL), which were grown for 3 hours before being induced with IPTG to a final concentration of 0.5 mM. For determination of holo-SOD1 variants, ZnSO_4_ and CuSO_4_ were added at induction to concentrations of 200 µM and 3 mM respectively. After induction, cells were grown for 4 hours and harvested (10,000*g* for 5 min) before being frozen at −80 °C until use. Cells were resuspended into lysis buffer (50 mM Tris, 500 mM NaCl, 1% Triton X-100, DNAse 20 µg/mL, 1 mM TCEP, 1 cOmplete™ EDTA-free protease inhibitor tablet (Sigma-Aldrich, USA) per 50 mL buffer (pH 7.4) and subject to 5 rounds of freeze/thawing at −80 °C and 42 °C. Following lysis, insoluble material was pelleted (20,000*g* for 5 min) and the supernatant collected as the soluble fraction. The insoluble pellet was resuspended in lysis buffer and repelleted (20,000*g* for 5 min) before being resuspended again in solubilizing buffer (7 M Urea, 2 M Thiourea, 25 mM TCEP, 4% CHAPS (w/v)) and incubated at room temperature with gentle rotation for 1 hour. After solubilisation, samples were spun to clear any residual insoluble material (20,000*g* for 10 min) and the solubilized material was collected as the urea soluble fraction. Both the soluble and urea soluble fraction were diluted to equal volumes and were mixed with reducing LDS-sample buffer before being run on SDS-PAGE and stained with coomassie blue to determine the relative amounts of SOD1 in each fraction via densitometry. Measurements were made using FIJI(24).

### Mammalian Tissue Culture

U2OS cells (ATCC HTB-96) were cultured in Advanced Dulbecco’s modified Eagles medium-F12 (Adv. DMEM-F12) (Invitrogen, USA), supplemented with 10% (v/v) heat inactivated fetal bovine serum (FBS) (ThermoFisher, USA) and 2 mM GlutaMAX (ThermoFisher, USA). NSC-34 cells(25) were cultured in Adv. DMEM-F12 with 2% FBS and 2 mM GlutaMAX (ThermoFisher, USA). In order to passage and plate cells, they were washed once with pre-warmed 1× PBS and treated with 0.25% trypsin, 0.02% EDTA dissociation reagent (Invitrogen, USA) to lift off the adherent cells. The cells were pelleted via centrifugation (300*g* for 5 min) and resuspended in pre-warmed culture media. Following washing, plates, cover slips and chamber-slides were seeded at a confluency of 40% and cultured at 37 °C in a humidified incubator with 5% atmospheric CO_2_ for 24 h prior to transfection (~70-80% confluent). Cells were transfected with plasmid (0.5 µg per well of a 24-well plate and 0.25 µg per chamber of an 8-well chamber slide unless state otherwise) 24 h post-plating using Lipofectamine LTX (Invitrogen, USA) according to the manufacturer’s instructions.

### Imaging of Fixed Cells

U2OS cells (selected for their flat morphology, ease of transfection, and easily measurable inclusion phenotype) transfected with SOD1-EGFP plasmids were prepared for imaging 24 hr post-transfection using the following methods. Cells were washed once with pre-warmed (37 °C) 1× PBS and incubated in 4% paraformaldehyde (PFA) in PBS for 20 min. Fixed cells were then washed twice with 1 × PBS (5 min per wash), and then permeabilized using 0.1% Triton X-100 in PBS (5 min). Permeabilized cells were washed twice with 1× PBS (5 min per wash) and nuclei were stained with Hoescht 33342 (1 µg/mL in 1× PBS) for 5 min. Following staining of nuclei, cells were washed twice in 1 × PBS (5 min per wash) and either imaged immediately or stored at 4 °C until imaging (no longer than 48 h).

Images were acquired using an inverted Axio Observer Z1 microscope (Carl Zeiss AG, Germany) equipped with an AxioCam HighRes camera (Carl Zeiss AG, Germany) and a motorized stage. An A-Plan 10×/0.25NA air objective was used to capture images with a 62 HE BFP/GFP/HcRED reflector, a 395-495-610 beam splitter, with a Colibri LED light source (Carl Zeiss AG, Germany) where Hoescht 33342 (45 ms exposure time - 25% light source intensity) was excited using 350 – 390 nm light and its emission was captured at 402 – 448 nm, and GFP (50 ms exposure time - 50% light source intensity) was excited with 460 – 488 nm light and its emission captured from 500 – 557 nm. Camera settings were 2× analogue gain with 1,1 binning mode acquiring 16 bit images at 1388 × 1040 pixel resolution in a 3×3 grid for adequate well coverage. Importantly, these image acquisition settings were optimized to minimize saturation of GFP signal so that inclusions would be accurately determined using analysis algorithms. The microscope was controlled with Zen 2.5 Pro software (Carl Zeiss AG, Germany).

### Live-Cell Imaging

NSC-34 cells were plated into Lab-Tek 8-well chamber slides (Thermofisher, USA) and transfected with SOD1-EGFP constructs as described above 24 h after plating. Transfected cells were changed into imaging media (Fluorobrite™ DMEM, 25 mM HEPES, 2 % FBS) with 3 µM DRAQ7 (Abcam, UK) 24 h post-transfection and were allowed to equilibrate on a heated microscope stage to 37 °C with 5 % atmospheric CO_2_ for 30 min before imaging began. Images were acquired using an EC Plan-Neofluar 5×/0.17NA M27 air objective every 30 min. EGFP was was excited similarly as above except for a shorter exposure time of 10 ms was used whereas DRAQ7 was excited with 562-607 nm light (100 ms exposure) and its emission captured from 615-800 nm. Images were binned 2×2 to reduce phototoxicity and bleaching (15 % blue light source intensity and 50% yellow light source intensity). Focus was maintained using a definite focus module. These settings were kept consistent between replicates and cells were imaged for 72 h.

### Analysis of Inclusion Formation in Cultured Cells

Image analysis of transfected U2OS cells was carried out using a combination of Zen 2 lite (Carl Zeiss AG, Germany), CellProfiler (version 3.0) and CellProfiler Analyst (version 2.2.1) (4). Separate channels (Hoescht 33342 and GFP) were exported from Zen 2 lite in 16-bit PNG image format without compression. Images were then imported into CellProfiler and examined using a custom pipeline designed to identify and segment cells (see SI Appendix SFigs 2,3 for example images of this process, and Supplementary File of the pipeline for download). The pipeline consisted of three main parts: The first processes were optimized to identify and segment nuclei in the Hoescht 33342 channel, the second process was optimized to identify and segment the cell cytoplasm for determination of transfected cells, and the third process was optimized to measure parameters of each individual cell for downstream assisted machine learning analysis. At each stage of analysis, the actions of the pipeline were visually inspected to ensure proper segmentation and measurement.

Nuclei were first smoothed with a Gaussian filter (artifact diameter 4 pixels) to remove local deviations of fluorescence intensity within each nuclei, followed by which background signal was subtracted using a top-hat transform method (SI Appendix SFig 3). Nuclei were then identified and segmented based upon their diameter and fluorescence intensity. Following identification, nuclei were shrunk to improve the accuracy of cytoplasmic identification and segmentation. The cell cytoplasm was determined and segmented in the GFP channel using a propagative method with adaptive Otsu thresholding (SI Appendix SFig. 2). Untransfected cells were filtered out by selecting only those nuclei which had an overlapping GFP signal above background.

Once the proper selection and segmentation of cells from images was visually confirmed, three key properties were measured for the determination of cells with inclusions *vs*. those without: Mean GFP intensity, intensity distribution, and texture. Mean GFP intensity was used because cells with greater levels of SOD1 expression typically have more inclusions(26). Misfolded SOD1 tends to be partitioned to the juxtanuclear quality control compartment (JUNQ) which is proximal to the nucleus of a cell(27). We thus measured the intensity distribution of EGFP signal within individual cells. Cells with inclusions were expected to have intensity values that were initially higher closer to the cell centre, and to exhibit a sharp decline towards the cyotplasmic periphery; Cells with no inclusions would have a smoother decline in fluorescence signal from the middle to the periphery edge. Lastly, we measured the texture (intensity variations) of the GFP signal within individual cells, since we expected cells with inclusions to have much higher texture values due to the inclusions being much brighter in local areas than the diffuse and evenly distributed cytosolic EGFP signal.

Following analysis of images with CellProfiler, an exported SQL database containing location and measurement data for each individual cell was loaded into CellProfiler Analyst for assisted machine learning. The entire dataset analysed included 890 total images from three biological replicates of each SOD1-EGFP mutant, with each image containing on average 200 (~ 100 transfected) cells for analysis. We categorized cells as negative (no inclusions), positive (inclusions – puncta within cells that was brighter than cytoplasmic background), or dead (small and rounded, and/or blebbing morphology) based upon expert visual inspection of individual cells. The trainer was blinded to which construct was transfected into the individual cells to remove bias from the analysis. A random forest classifier was trained on 304 positive, 337 negative and 81 dead cells, and was verified by requesting cells from each category until ≥ 95% of cells chosen by the classifier were determined to be correct by the viewer. Training was also confirmed by asking the classifier to score random images with known values. Following verification of correct training, the classifier was used to score all images to classify each individual cell within each image. The image dataset contained approximately 90,000 unique transfected cells, with approximately 7500 transfected cells analysed per transfected construct. For co-transfection experiments, a random forest classifier was trained on 408 positive, 527 negative and 125 dead cells of an image dataset containing ~50 000 unique transfected cells. The CellProfiler pipeline is available as a supplementary file.

### Analysis of Live-Cell Imaging

Images were exported from Zen 2 lite (Carl Zeiss, Germany) as 16-bit PNG files with original data. DRAQ7 images were pre-processed with ImageJ using the ‘triangle’ auto threshold method to account for the moving background signal, which resulted in a binary image. Images were then imported into CellProfiler for analysis using a custom pipeline designed to identify and count transfected cells that were also DRAQ7 positive. Briefly, GFP images were histogram equalized to make fluorescence values within and between images equal, then background was subtracted using the *enhance features* module with a top hat transform. GFP positive cells were identified using the *identify primary objects* module based on size and fluorescence intensity and were counted across image sets. DRAQ7 positive cells were identified using the *identify primary objects* module from the binary images. The *relate objects module* was used to identify which GFP positive cells were also DRAQ7 positive. The raw number values were normalized to the starting number of cells so that proliferation could be compared between replicates.

### Zebrafish ethics statement

Fish care and experimental protocols were approved under the protocol AUP00000077 by the Animal Care and Use Committee: Biosciences at the University of Alberta under the auspices of the Canadian Council on Animal Care.

### A89R and K128N mutagenesis and mRNA synthesis

Plasmids containing the coding sequences for human wildtype, A4V, and G127X SOD1 variants were cloned into the pCS2+ plasmid via Gateway cloning as previously described(28). The *pCS2+.SOD1^WT^* plasmid was utilized for mutagenesis reactions to create the novel A89R and K128N mutations. Production of mCherry and Tol2 transposase mRNA was also performed using pCS2+ plasmids containing these coding sequences.

Mutagenesis was performed using the Agilent QuikChange Lightning Site-Directed Mutagenesis kit (Agilent Technologies, Cat. No. 210518), with primers designed using the online Agilent primer design program (http://www.genomics.agilent.com/primerDesignProgram.jsp): For K128N, forward primer 5’-aagcagatgacttgggc<u>aa**c**</u>ggtggaaatgaagaaag- 3’ and reverse primer 5’-ctttcttcatttccacc<u>**g**tt</u>gcccaagtcatctgctt-3’; for A89R, forward primer 5’-tgttggagacttgggcaatgtgact<u>**cg**t</u>gacaaagatggt-3’ and reverse primer 5’-accatctttgtca**cg**agtcacattgcccaagtctccaaca-3’ (letters in bold indicate mutagenesis sites, and underlined triplets indicate the amino acid codon). Mutagenized plasmid was then transformed into One Shot Top10 chemically competent cells (Invitrogen, Cat. No. C4040-03). Colonies were sequenced to confirm mutagenesis and stored as glycerol stocks. SOD1 variants, mCherry, and inert transposase mRNA for injection were synthesized by linearizing pCS2+ plasmids with FastDigest NotI restriction enzyme (Thermo Fisher Scientific, Cat. No. FD0593), and performing transcription reactions with the mMESSAGE mMACHINE SP6 transcription kit (Ambion, Cat. No. AM1340). After transcription, mRNA products were verified using gel electrophoresis with a RiboRuler High Range ladder, prepared according to package instructions (Thermo Scientific, Cat No. SM1821), then aliquoted and stored at - 80°C until use.

### Animal care and embryo injections

Zebrafish were raised and maintained under standard procedures (29). Embryos were collected and kept in E3 embryo media at 28°C with PTU added at 6-8 hpf to prevent pigmentation. The *Tg(mnx1:GFP)* transgenic line (ZFIN ID: ZDB-ALT-051025-4) (30, 31) was utilized for visualization of the primary motor axons via GFP fluorescence. Adult transgenic fish were crossed to wild type AB fish to maintain consistency of GFP expression among embryos. Embryos were injected into yolk at the 1-2 cell stage with a total of 1900 pg of mRNA in the combinations listed in Supplementary Table 3, and screened at 24 hpf for mCherry fluorescence and GFP expression in the spinal cord. The SOD1 mRNA dosages used here were within ranges used in previous studies(32), to establish an axonopathy phenotype detectable above control levels. Inert transposase mRNA was utilized as a “top-up” to ensure consistent 1900 pg total dose among all groups.

### Assessment of Zebrafish Motor Axons

At 33-37hpf, embryos were fixed in 4% paraformaldehyde/5% sucrose/0.1M sodium phosphate for 40 minutes. Fixative was replaced with PBST and embryos stored at 4°C until primary motor axons could be assessed (assessments were completed within 24 hours of fixation). Assessments were performed manually on a Leica MZ16F stereo microscope with a PLANAPO 1.6× objective (10447050, Leica Microsystems) and a GFP2 Ultra filter set (10447407, Leica Microsystems). The primary motor axons were viewed by a blinded researcher and scored as normal or abnormal as previously described (32). Briefly, primary motor axons with branching occurring dorsal of or level with the ventral edge of the notochord were scored as abnormal (Fig. 5a, b main text). Motor axons exiting both lateral sides of the spinal cord were assessed. The total number of abnormal axons was recorded for each individual embryo and these values were averaged for each injection group. Treatment group *n*-values are: mRNA control - 68, K128N - 66, WT - 45, G127X - 38, A4V (900pg) - 55, A89R - 49, A4V (1800pg)- 83, A4V/WT −141, A4V/K128N - 101. For statistical analysis, Kruskall-Wallis tests with post-hoc Mann-Whitney pairwise comparisons were performed in PastProject for Mac, version 3.09 (Øyvind Hammer, University of Oslo, http://folk.uio.no/ohammer/past/). Representative images in Fig. 4 Main Text are for illustrative purposes only, as axonopathy assessments were performed manually as described above. Motor neuron imaging for Fig. 4 Main Text was performed on a Zeiss Axio Observer.Z1 microscope with 20× objective lens, LSM 700 confocal scanner, and Zen software (2010, Carl Zeiss Imaging). Transgenic embryos were mounted on slides in glycerol. Images were acquired in the 488 channel (1024 × 1024 pixels, 12 bit, gain set at 350-375). For images shown in Fig. 5 Main Text, z-stacks of motor neurons were flattened and adjusted for image rotation (for best view of axons), brightness, and contrast in Imaris x64 (version 7.4.0, Bitplane, Badnerstrasse).

### Quantification and Statistical Analysis

Statistical parameters are described in figure legends and/or the material and methods section, including description of error bars, sample replication numbers, p-values, as well as specific statistical analyses used.

### Data and Materials Availability

The data and materials that support the findings of this study can be made available upon request from the corresponding author.

**Fig. S1.**
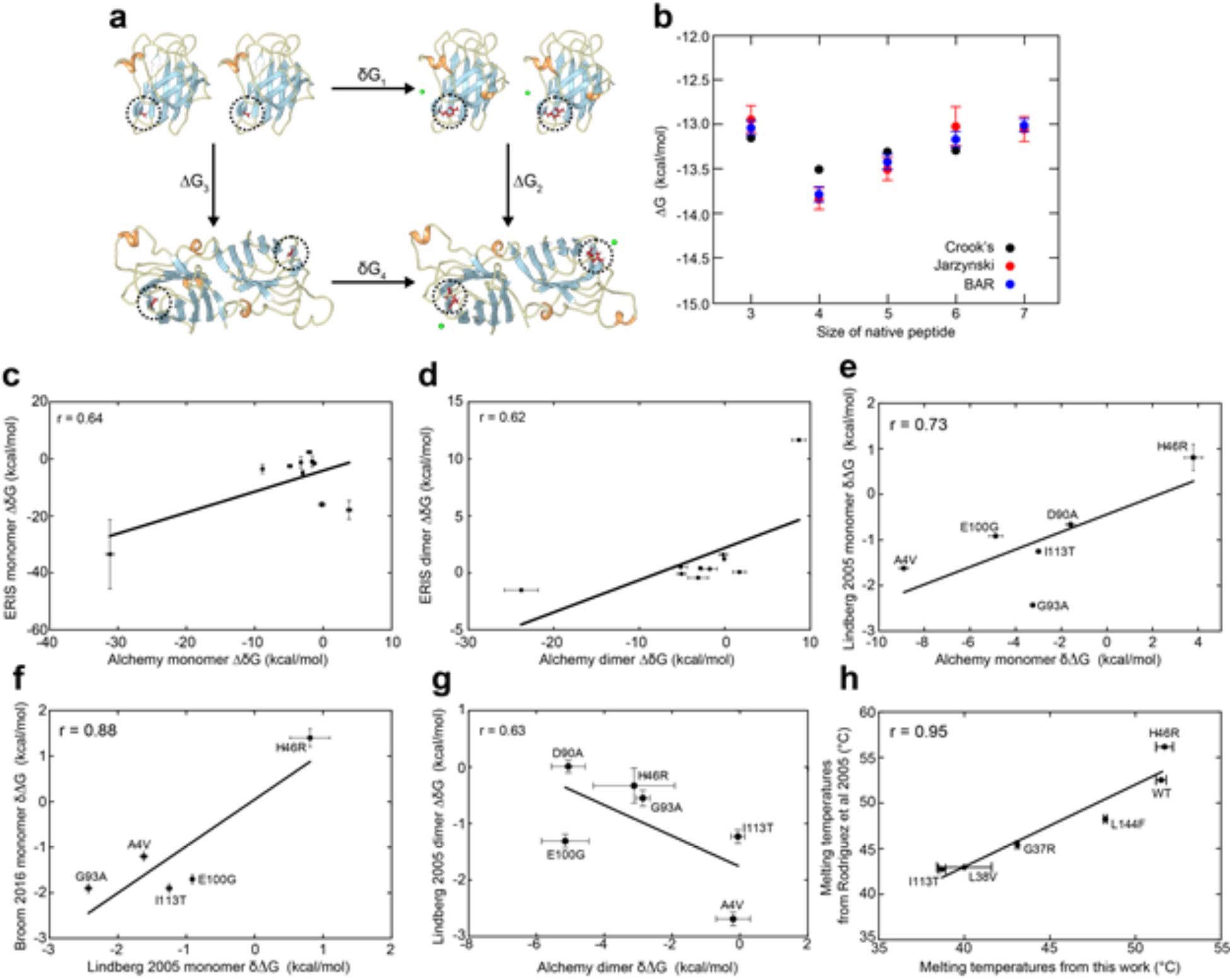
Comparison of values determined in this work to those in literature. **(a)** The dimer thermodynamic cycle when mutations (A89R in this case, circled licorice) are made to both SOD1 chains. A chloride ion must also be added (green) to preserve system neutrality. The thermodynamic cycle mandates that δ∆*G* = ∆*G_2_* - ∆*G_3_* = ∆*G_4_* - ∆*G_1_*. Alchemical computations of ∆*G_4_* and ∆*G_1_* thus determine the mutational change of dimer binding free energy, δ∆*G*. **(b)** Three separate methods of determining the alchemical free energy change upon mutation (Crooks theorem, Jarzynski’s theorem, and the Bennett acceptance ratio method (BAR)) are all in good agreement, and show good convergence as the length of the unfolded peptide (centered on the residue of interest) is increased. **(c)** Correlations of the free energy difference of monomer unfolding between ERIS server predictions and alchemical modelling. (r = 0.64, p = 0.0939). **(d)** Correlation of the free energy difference of dimer binding between ERIS server predictions and alchemical modelling. (r = 0.62, p = 0.0451). The modest correlation in (c,d) indicates that the more intensive alchemical method is warranted. **(e)** Comparison of alchemically derived monomer unfolding free energy *vs.* unfolding free energy determined in Lindberg (2005)(1) (r = 0.73, p = 0.0953). **(f)** Correlation of monomer unfolding free energies determined in Lindberg (2005)(1) *vs.* those in Broom (2016)(2) (r = 0.88, p = 0.0519). **(g)** Correlation of dimer binding free energy from alchemical modelling and those determined in Lindberg (2005)(1) (r = 0.63, p = 0.1796). **(h)** Correlation of melting temperatures determined by DSF in this work with those determined by DSC in Rodriguez (2005)(3) (r = 0.95, p = 0.0032). For both the previous data and the present study, E,E(SS) WT SOD1 was used as the background for mutations.

**Fig S2.**
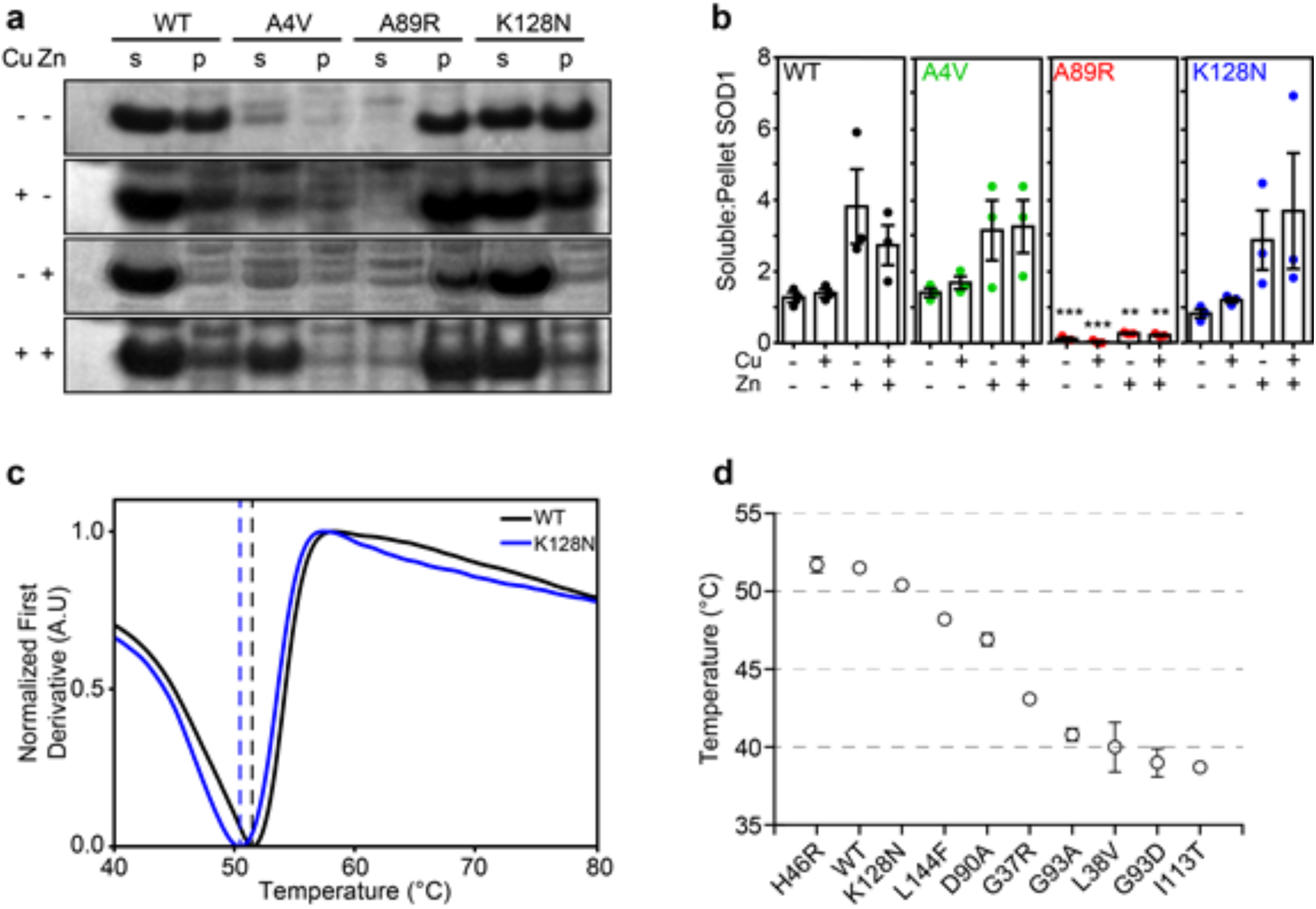
A89R forms insoluble inclusions when expressed in bacterial cells regardless of metalation state, whereas K128N has WT-like solubility and stability. **(a)** Cell lysates containing soluble (s) and insoluble pellet (p) fractions of WT, A4V, A89R, and K128N SOD1, expressed in bacteria with or without Cu and/or Zn. Uncropped gels are shown in supplementary information. **(b)** Quantification of the soluble and insoluble fractions of bacterially expressed SOD1 showing that A89R is insoluble even in the presence of metal cofactors; the solubility of K128N shows no significant difference from WT. **(c)** First derivatives of DSF melting curves for SOD1 WT and K128N show similar melting temperatures for the two mutants. **(d)** Rank ordered average T_m_’s of all SOD1 mutants purified in this study (see supplementary table 2 for details). Error bars represent SD of the mean. Significance was determined by one-way ANOVA with Tukey’s post-test where mutants were compared to WT for their respective treatments (*** *p* < 0.01, ** *p* < 0.05).

**Fig. S3.**
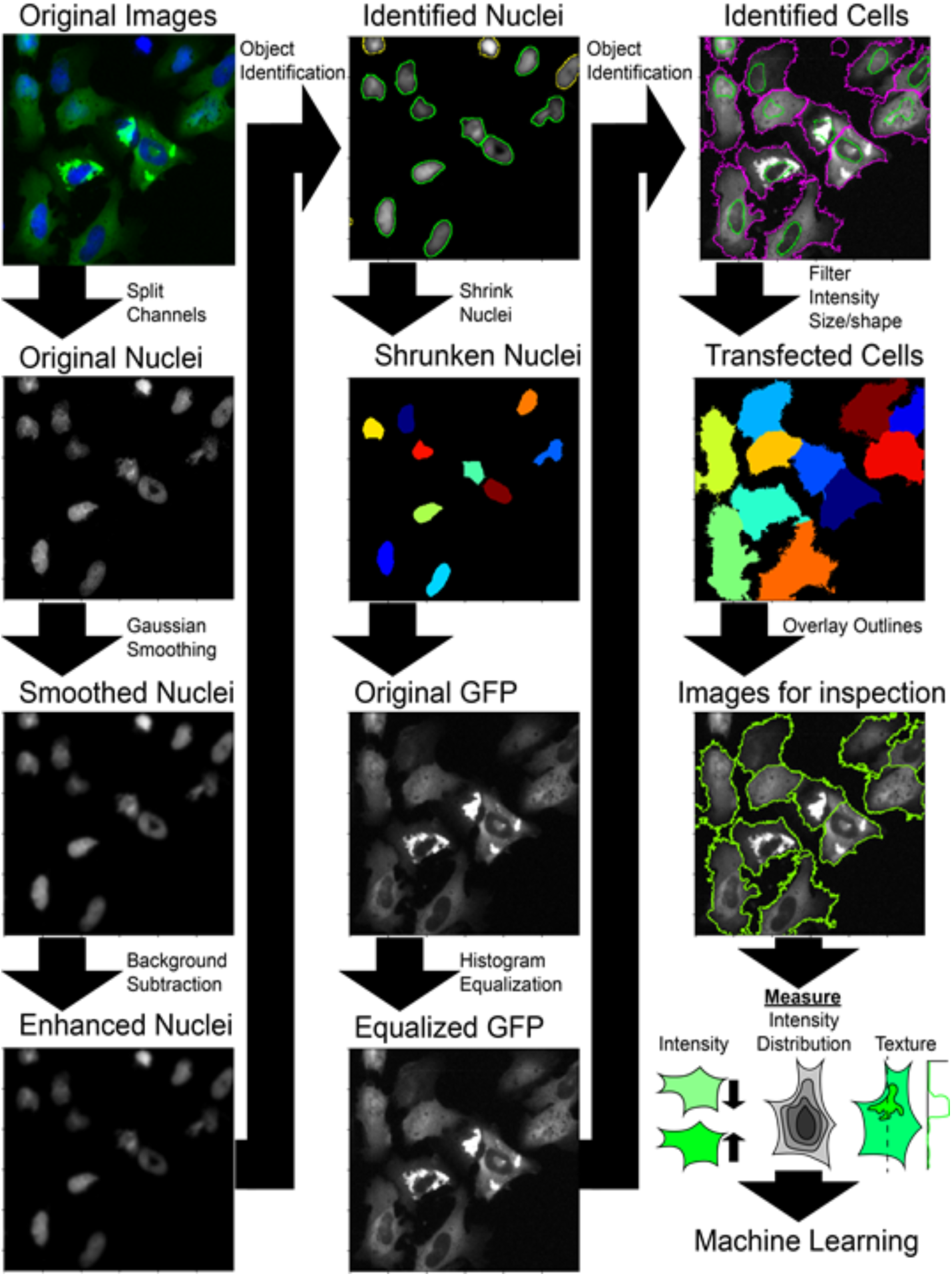
Example of image processing and measurement for assisted machine learning. Images are acquired in the blue (nuclei) and green (GFP) channel. Following acquisition, channels are exported as separate images and loaded into CellProfiler(4). Firstly, images are preprocessed before measurement to obtain accurate segmentation. Nuclei are smoothed to remove local noise that may result in a single nuclei being split into two, they are then enhanced by subtracting background with a top-hat rolling ball method. Identification is performed on background-subtracted nuclei based upon size and intensity. The nuclei are then shrunk to make segmentation of cytoplasm easier. GFP images are then processed by histogram equalization with a cell-sized grid to equalize the fluorescence intensity across all transfected cells. The cytoplasmic cell boundary is then identified, and non-transfected cells filtered out by measuring their size and fluorescence intensity. Transfected cell outlines are then overlaid on each image and exported for visual inspection prior to measurement. Following verification of correct cell identification, the mean intensity, texture (intensity variance), and spatial intensity distribution within individual cells are measured as key values for assisted machine learning. Data and images are exported and loaded into CellProfiler Analyst for training and analysis.

**Fig. S4.**
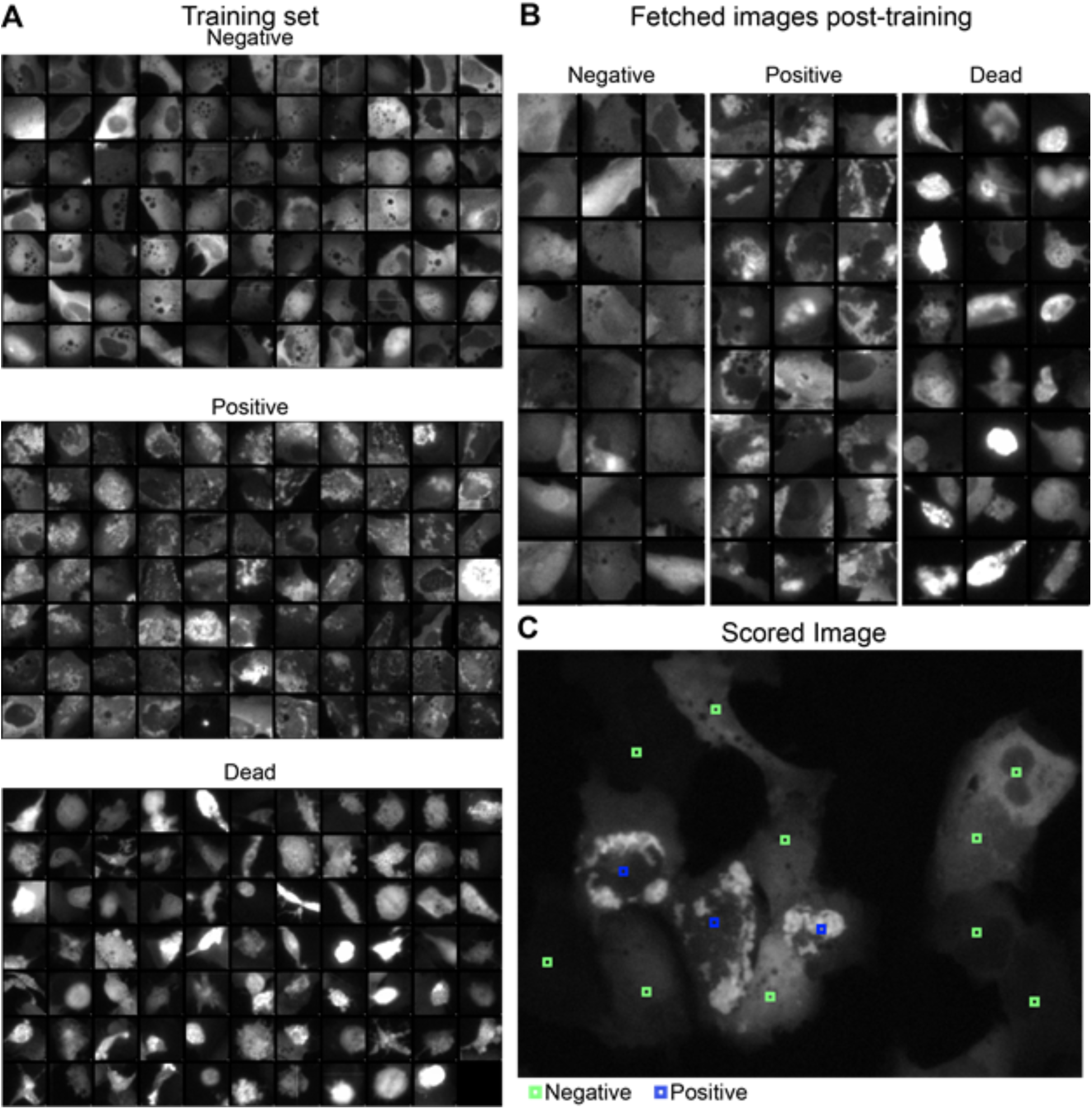
Example of machine learning inputs for identification of cells with inclusions. **(A)** Blinded thumbnail images of cells are classified by visual inspection into 3 categories: negative (cells without inclusions), positive (cells with inclusions), and dead (blebbing, rounded, small). **(B)** After adequate classification, the algorithm is trained and cells are fetched from the pool of cell not used for training, for verification of accuracy. **(C)** Once the algorithm is deemed accurate at determining cells with inclusions (≥ 95% fetched cells scored correctly in each group), the entire image set is scored.

**Supplementary Table 1.** List of SOD1 variants used in this work and corresponding experiments.

| **SOD1 Variant** | **Alchemical Modelling** | **Differential Scanning Fluorimetry** | **Bacterial Solubility** | **U2OS inclusion formation** | **NSC-34 toxicity Assay** | ***Danio rerio* Injections** |
| --- | --- | --- | --- | --- | --- | --- |
| **Wild type** | + | + | + | + | + | + |
| **A4V** | + | - | + | + | + | + |
| **G37R** | + | + | - | + | - | - |
| **L38V** | - | + | - | + | - | - |
| **H46R** | + | + | - | + | - | - |
| **A89R** | + | - | + | + | + | + |
| **D90A** | + | + | - | + | - | - |
| **G93A** | + | + | - | + | - | - |
| **G93D** | + | + | - | + | - | - |
| **E100G** | + | - | - | - | - | - |
| **I113T** | + | + | - | + | - | - |
| **K128N** | + | + | + | + | + | + |
| **G127X** | - | - | - | - | - | + |
| **L144F** | - | + | - | + | - | - |

| **SOD1 Variant** | **Alchemical Modelling** | **Differential Scanning Fluorimetry** | **Bacterial Solubility** | **U2OS inclusion formation** | **NSC-34 toxicity Assay** | ***Danio rerio* Injections** |
| --- | --- | --- | --- | --- | --- | --- |
| Wild type | + | + | + | + | + | + |
| A4V | + | - | + | + | + | + |
| G37R | + | + | - | + | - | - |
| L38V | - | + | - | + | - | - |
| H46R | + | + | - | + | - | - |
| A89R | + | - | + | + | + | + |
| D90A | + | + | - | + | - | - |
| G93A | + | + | - | + | - | - |
| G93D | + | + | - | + | - | - |
| E100G | + | - | - | - | - | - |
| I113T | + | + | - | + | - | - |
| K128N | + | + | + | + | + | + |
| G127X | - | - | - | - | - | + |
| L144F | - | + | - | + | - | - |

**Supplementary Table 2.** Melting points of apo-SOD1 variants as determined by differential scanning fluorimetry.

| **SOD1 Variant** | **T<sub>m</sub> (°C)** | **ΔT<sub>m</sub>(°C)** |
| --- | --- | --- |
| Wild-type | 51.5 ± 0.3 | - |
| G37R | 43.1 ± 0.1 | -8.4 ± 0.3 |
| L38V | 40.0 ± 1.6 | -11.5 ± 1.6 |
| H46R | 51.7 ± 0.5 | 0.2 ± 0.6 |
| D90A | 46.9 ± 0.4 | -4.6 ± 0.5 |
| G93A | 40.8 ± 0.4 | -10.7 ± 0.5 |
| G93D | 39.0 ± 0.9 | -12.5 ± 0.9 |
| I113T | 38.7 ± 0.2 | -12.8 ± 0.4 |
| K128N | 50.4 ± 0.2 | -1.1 ± 0.4 |
| L144F | 48.2 ± 0.1 | -3.3 ± 0.3 |

**Supplementary Table 3.** List of injections and corresponding mRNA amounts for zebrafish axonopathy experiments. Picograms = pg.

| **Treatment Group** | **SOD1 Variant mRNA (pg)** | **mCherry mRNA (pg)** | **Tol2 mRNA (pg)** | **Total mRNA (pg)** | **n (Fish)** |
| --- | --- | --- | --- | --- | --- |
| **mRNA Control** | *None* | 100 | 1800 | 1900 | 68 |
| **K128N 900** | K128N 900 | 100 | 900 | 1900 | 66 |
| **WT 900** | WT 900 | 100 | 900 | 1900 | 44 |
| **G127X 900** | G127X 900 | 100 | 900 | 1900 | 38 |
| **A4V 900** | A4V 900 | 100 | 900 | 1900 | 55 |
| **A89R 900** | A89R 900 | 100 | 900 | 1900 | 49 |
| **A4V 1800** | A4V 1800 | 100 | 0 | 1900 | 84 |
| **A4V 900 + WT 900** | A4V 900<br>WT 900 | 100 | 0 | 1900 | 63 |
| **A4V 900 + K128N 900** | A4V 900<br>K128N 900 | 100 | 0 | 1900 | 101 |

